## Supplemental Table 1 for "The holocephalan ratfish endoskeleton shares trabecular and areolar mineralization patterns, but not tesserae, with elasmobranchs little skate and catshark"

**Table 2. Measurements and regions of interest of specimens**

| Specimens | Sample size | Total length TL and disc width DW measurements | Regions of interest |
|---|---|---|---|
| Little skate stage 32 embryos | 5 | 3.2 cm DW, 7.5 cm TL<br>2.8 cm DW, 7 cm TL<br>2.8 cm DW, 7.3 cm TL<br>3 cm DW, 7.3 cm TL<br>2.9 cm DW, 7.3 cm TL | Caudal vertebrae |
| Little skate stage 33 embryos | 5 | 3.5 cm DW, 8 cm TL<br>3.6 cm DW, 7.8 cm TL<br>3.4 cm DW, 7.8 cm TL<br>3.4 cm DW, 7.5 cm TL<br>3.3 cm DW, 7.9 cm TL | Caudal vertebrae |
| Little skate juveniles | 5 | 5.5 cm DW, 10.4 cm TL<br>5.6 cm DW, 10 cm TL<br>6 cm DW, 10.5 cm TL<br>6 cm DW, 11 cm TL<br>6.5 cm DW, 11 cm TL | Caudal vertebrae |
| Little skate adults | 4 | 43.5 cm TL<br>45 cm TL<br>47 cm TL<br>47.5 cm TL | Precaudal vertebrae, caudal vertebrae |
| Small-spotted catshark | 3 | 31 cm TL<br>31 cm TL<br>31 cm TL | Ceratohyal, precaudal vertebrae, caudal vertebrae |
| Spotted ratfish | 5 | 28.5 cm TL<br>32 cm TL<br>33 cm TL<br>40 cm TL<br>45 cm TL | Precaudal vertebrae, pharyngeal skeleton, ceratohyal, and synarcual |
